# Intraspecific variation in thermal tolerance differs between tropical and temperate fishes

**DOI:** 10.1101/2020.12.07.414318

**Authors:** J.J.H. Nati, M.B.S. Svendsen, S. Marras, S.S. Killen, J.F. Steffensen, D.J. McKenzie, P. Domenici

## Abstract

How ectothermic animals will cope with global warming, especially more frequent and intense heatwaves, is a critical determinant of the ecological impacts of climate change. There has been extensive study of upper thermal tolerance limits among fish species but how intraspecific variation in tolerance may be affected by habitat characteristics and evolutionary history has not been considered. Intraspecific variation is a primary determinant of species vulnerability to climate change, with implications for global patterns of impacts of ongoing warming. Using published critical thermal maximum (CT_max_) data on 203 marine and freshwater fish species, we found that intraspecific vsariation in upper thermal tolerance varies according to a species’ latitude and evolutionary history. Notably, freshwater tropical species have lower variation in tolerance than temperate species in the northern hemisphere, which implies increased vulnerability to impacts of thermal stress. The extent of variation in CT_max_ among fish species has a strong phylogenetic signal, which may indicate a constraint on evolvability to rising temperatures in tropical fishes. That is, in addition to living closer to their upper thermal limits, tropical species may have higher sensitivity and lower adaptability to global warming compared to temperate counterparts. This is evidence that tropical fish communities, worldwide, are especially vulnerable to ongoing climate change.

---

The capacity of ectothermic species to cope with ongoing global warming, especially the increasing frequency, intensity and duration of extreme heatwaves, will be influenced by their upper thermal tolerance limits ^1–3^. Tolerance of acute warming, measured as the critical thermal maximum (CT_max_), varies among fish species according to thermal conditions in their habitat ^4^. Tropical species live in warm, relatively thermally stable habitats; they have narrow thermal tolerance ranges but higher CT_max_ than species at temperate latitudes. Their warm habitat temperatures are also, however, closer to their limits of upper thermal tolerance, so they have a limited thermal safety margin (defined as the difference between upper thermal tolerance limit CT_max_ of adult life stage and the maximum habitat temperature during summer ^5^) and consequently are considered to be especially vulnerable to global warming ^6-9^. Temperate species have lower absolute thresholds for tolerance of warming, but they have broader tolerance ranges, presumably because they encounter a wide range of habitat temperatures, both seasonally and spatially. This is linked to wider thermal safety margins than in tropical species^4,10^. These patterns of vulnerability to global warming, among species at a geographic scale, are major issues in projecting impacts of warming. They have a strong phylogenetic basis, which is believed to reflect local adaptation to common ancestral thermal regimes in related species ^11^.

Studies of broadscale geographic patterns in vulnerability have, to date, focused upon average values for CT_max_ among fish species. The significance of intraspecific variation in tolerance remains to be explored. The extent of variation in functional traits within species, particularly of physiological tolerances, is expected to have a profound influence on their vulnerability to global change ^12-15^. Possessing a broad range of tolerance phenotypes in populations can reduce sensitivity to impacts of environmental stressors, through various proximate ecological mechanisms ^12-14^. If phenotypic variation is linked to underlying genetic diversity in the species, this can provide scope for adaptability and evolvability, by yielding genotypes for selection in changing environments ^12-14^. When fish species are challenged by thermal stressors such as increased seasonal temperatures and extreme heatwaves, the population sensitivity and adaptability will be major determinants of their relative vulnerability ^13-16^ (Figure 1).

**Figure 1.**
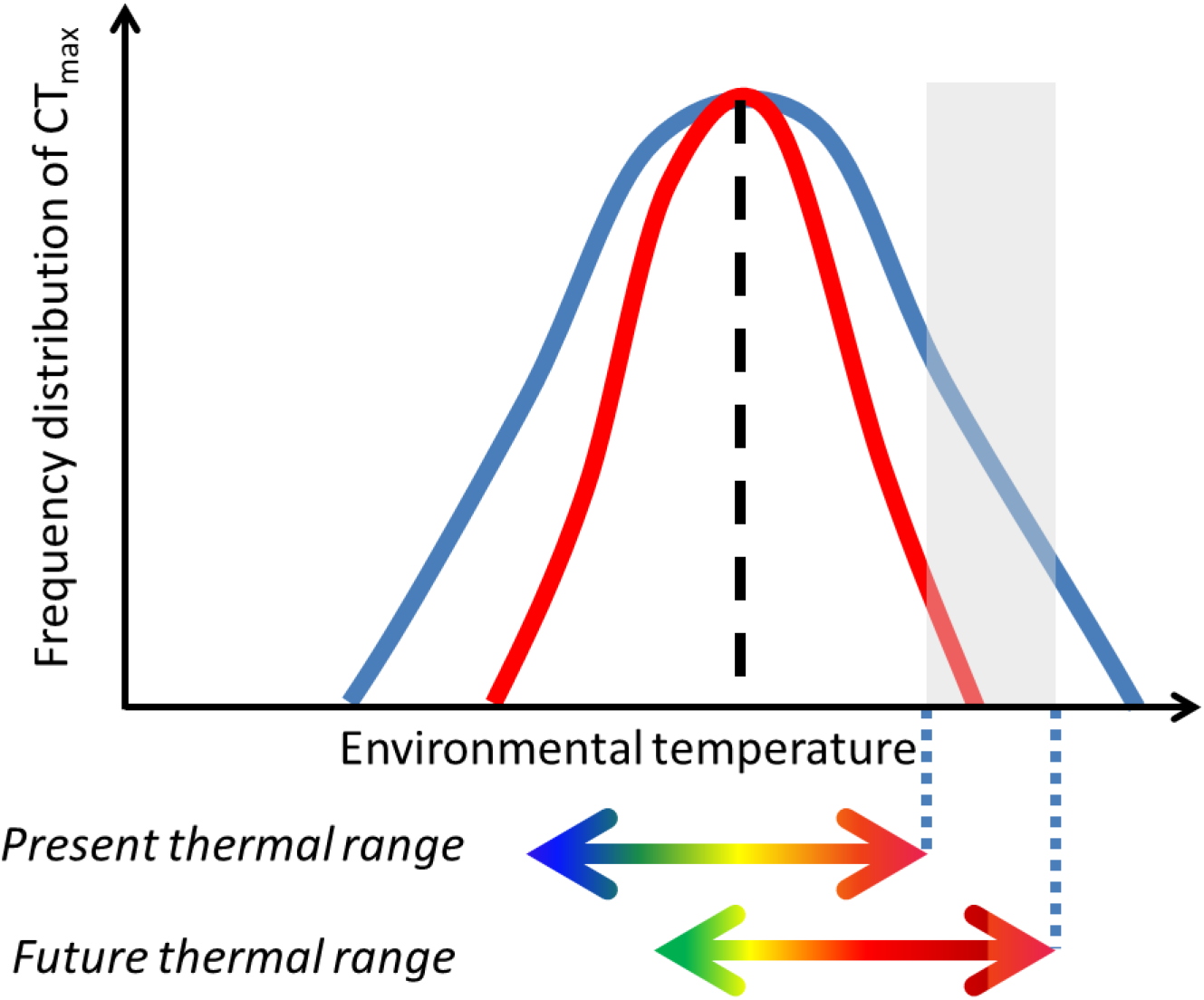
Theoretical representation of different frequency distribution curves of CT_max_. The curves of two species have the same mean CT_max_ (dashed line) but different standard deviations (S.D.). With ongoing climate change, represented by the shift in the thermal range (double-pointed arrows), individuals of the species with the narrower S.D.CT_max_ (red curve) are less likely to survive compared to individuals of the species with the wider S.D. CT_max_ (blue curve), since maximum enviromental temperatures will include values (grey area) outside their thermal tolerance range.

Fish species show intraspecific variation in CT_max_, which has a component of both phenotypic plasticity and heritable genetic variation ^15, 17-19^. The CT_max_ varies among populations of fish species, due to local adaptation ^20-22^, indicating that the trait evolves in response to prevailing thermal regimes. Given the broader thermal range experienced by temperate fish species, within generations and over evolutionary time, we hypothesized that they would exhibit greater intraspecific variation in their thermal tolerance, measured as CT_max_, than tropical species. We expected that the extent of variation might be linked to the magnitude of the thermal safety margin, because a small margin might constrain scope to express variation ^10^. We also expected the extent of variation in CT_max_ to have a phylogenetic basis, indicating that it reflected evolutionary processes of adaptation.

We used published data ^4^ and, after a data selection process (see Methods), we estimated the extent of intraspecific variation in CT_max_ of 203 species of ray-finned (actinopterygian) fish (n = 127 freshwater, n = 76 marine), based on the standard deviation of the mean. We were well aware that the selected studies in the dataset did not have the same protocol procedures. They did not use the same heating rate (0.0017-1°C/min) and fish size, both of which can influence the outcoming CT_max_ and standard deviation of the mean. We choose to not include these variables in our analysis because of the high variation of heating rate used and for fish size most studies did not report the size. We then compared two latitudinal groups, temperate to tropical species, considering the boundary to be 23° latitude. We also evaluated if variation in CT_max_ depended on whether species were from northern or southern hemisphere or whether species were marine or freshwater. Finally, we used the magnitude of the difference between acclimation temperature (T_a_) and CT_max_, which we denoted delta temperature (ΔT = CT_max_-T_a_), as an indication of the capacity to increase CT_max_ depending on the acclimation temperature, and evaluated if it was linked to intraspecific variation in CT_max_. All of the results were based on a phylogenetically informed analysis (phylogenetic least squares regression, PGLS, see Methods), to establish how patterns in the extent of variation were linked to evolutionary thermal history of the species.

There were significant differences in intraspecific variation in thermal tolerance in the two latitudinal groups (Figure 2). Freshwater tropical species showed lower intraspecific variation in CT_max_ (log_10_ S.D. CT_max_) than temperate (tropical species: PGLS, t = -2.054, p = 0.041, Figure 2). Species from northern hemisphere species had significantly lower variation in log_10_ S.D. CT_max_ than southern ones (PGLS, t = 2.318, p = 0.022; Figure 2A). Marine species did not differ from freshwater species (Figure 2B, PGLS, t = -1.683, p = 0.094). The ΔT had no significant association with log_10_ S.D. CT_max_ (PGLS, t = 1.972, p = 0.05; Figure S1.). There was no interaction between latitude and ΔT on log_10_ S.D. CT_max_. Phylogenetic relatedness among species contributed strongly to observed variation in log_10_ S.D. CT_max_ (PGLS, λ = 0.553, F_6,195_ = 4.397, p < 0.001, R^2^ = 11.92; Figure S3).

**Figure 2.**
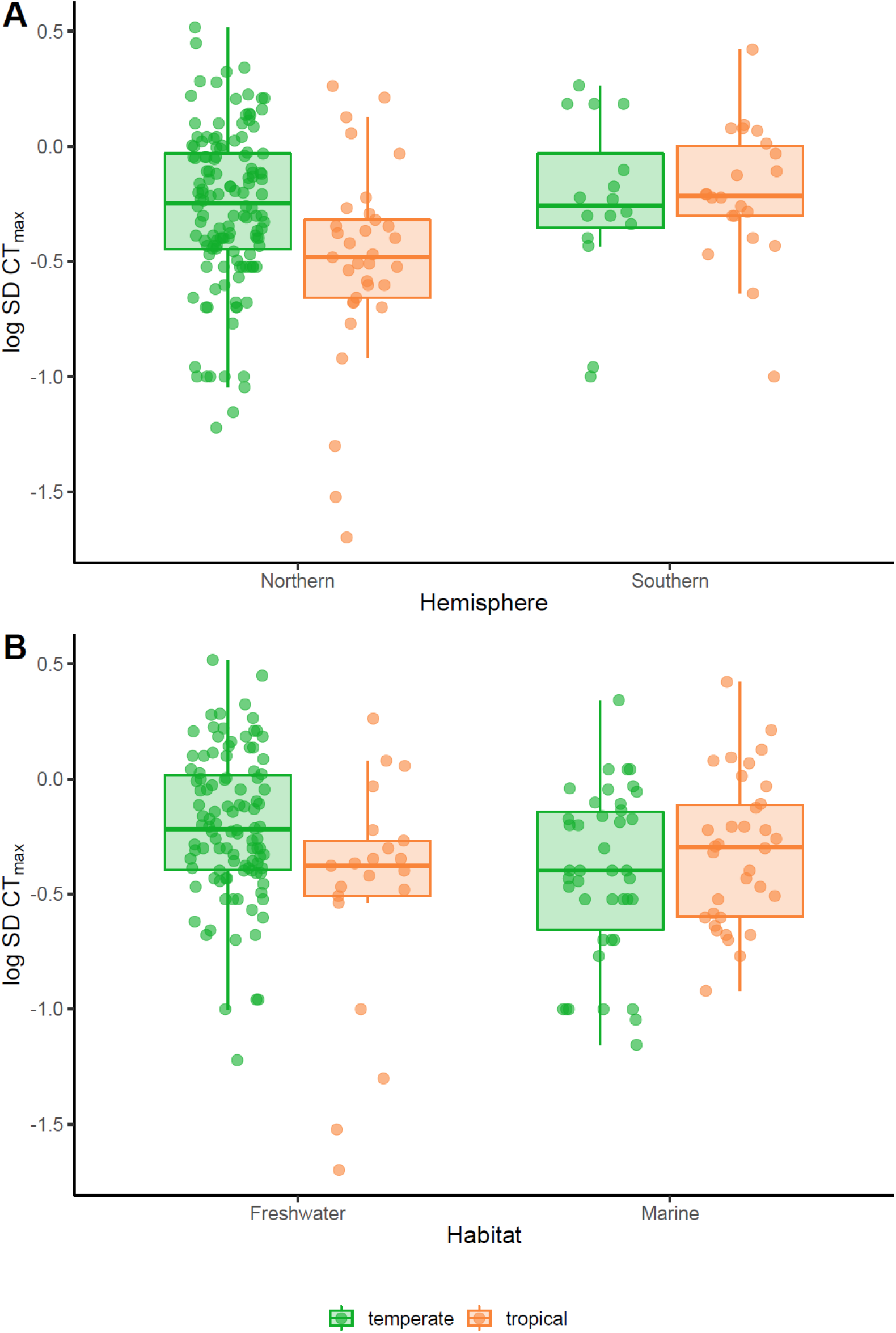
Intraspecific variation in CT_max_ (log_10_ transformed standard deviation CT_max_) divided into either temperate (148 species) or tropical (55 species). (**A**) Separated by hemisphere, Northern (132 temperate, 33 tropical species) or Southern (16 temperate and 22 tropical species). (**B**) Separated into freshwater (106 temperate, 21 tropical species) and marine (42 temperate, 34 tropical species).

These results show that freshwater tropical species have reduced within-species variation in thermal tolerance compared to temperate species. If this reflects a reduced capacity for phenotypic plasticity, this will increase their sensitivity to warming in the short term. If it reflects diminished heritable genetic variation, this will decrease adaptability and evolvability to a warmer and more thermally stressful future, over generational timescales. That is, the lower intraspecific variation in CT_max_ in freshwater tropical as compared to temperate species (Figure 2) renders the former especially vulnerable to future warming, in particular to extreme events ^24,25^ (Figure 1). This will compound the vulnerability of tropical species that derives from living near their upper thermal limits ^4,6,7,26^.

The fact that variation in thermal tolerance was more pronounced and variable in the northern compared to southern hemisphere could be the result of two phenomena: 1) greater thermal variability in the northern hemisphere ^4,6^; or 2) a relative paucity of data for the southern hemisphere ^27^. Nevertheless, the effect of hemisphere had a positive influence on intraspecific variation in CT_max_. Therefore, local thermal conditions experienced by species are determinant in setting the natural individual variation within populations.

The strong phylogenetic signal for the extent of intraspecific variation in CT_max_ is presumably because many families contain species with a relatively common history of thermal adaptation (see Figure S3). That is, they have occupied similar thermal regimes within temperate or tropical habitats. In particular, there is a latitudinal effect on family distributions, with some families only being present in temperate (e.g. Gadidae) or tropical (e.g. Apogonidae) habitats, although some cosmopolitan families have species in both (e.g. Gobiidea, Blennidae) (Figure S2). In addition to the geographic collinearity that may be occurring with some families, the phylogenetically based differences in intraspecific variation among species may cause evolutionary constraints on evolvability in the face of ongoing warming and exposure to extreme events. The extent of such constraints is not clear and would depend on the exact genes affecting thermal tolerance and how these are represented within each family ^11^. Further highlighting how temperature regime may shape evolutionary trajectories within closely related species or those with a common ancestor, with potential consequences for their vulnerability to thermal stress.

This evidence for higher vulnerability of tropical species to climate variability and extreme marine warming events ^28^ may have numerous ecological implications beyond simple tolerance thresholds. Tropical species may be obliged to seek thermal refugia in colder areas if these are available, potentially changing community structures ^9,29^; such distribution shifts could have major ecological consequences ^30,31^. Overall, the extent of intraspecific variation in CT_max_ must be considered in models that project impacts of warming on fishes. Intraspecific variation for tolerance in other environmental conditions such as hypoxia and acidification would be the next step for future research. Further research should focus on the mechanisms that underly latitudinal variation in CT_max_ and whether these reflect universal principles across all species.

## Methods

### Dataset and data selection process

We used the data on CT_max_ in marine, brackish and freshwater fish species (2722 observations unimputed data set) published by ^4^. We performed a three-step selection procedure to identify the species for this study. First, we excluded data where CT_max_ was measured using death as an endpoint (1256 observations) as these do not correspond to the accepted definition of CT_max_ (loss of equilibrium but not death) ^32^, so the temperatures recorded will have exceeded the critical threshold. Second, we excluded polar species because of the sample size (n = 5) and discarded brackish water species because no indication was given about the nature of the brackish habitat (e.g. lagoon, estuary or others). Third, several species were tested at different acclimation temperatures resulting in multiple CT_max_ measures for the same species. We therefore took CT_max_ values measured at the lowest or mid-point tested acclimation temperature with the largest sample size of individuals used. This data selection procedure produced a dataset of 203 fish species for which we have S.D. of their CT_max_ (standard deviation).

### Calculation of delta temperature

We calculated the ΔT

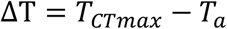

The ΔT defines the distance from thermal acclimation to thermal tolerance limit, providing an index of vulnerability to acute heating ^10^. This accounts for the fact that acclimation temperature is often asymptotically linked to CT_max_^15,24^.

### Data analysis

Analyses and models were made in R (3.4.4, R Foundation for Statistical Computing) using the phylogenetic generalized least squared method ^33,34^(PGLS, with caper package ^35^. Model selection was completed by AIC values using the AIC function estimating the best model fit (see Suppl. Table 1). The phylogeny of 203 fish species was found and generated from the comprehension tree of life (Suppl. Figure S3) ^36^ using the “rotl” package ^37^. A measure of phylogenetic correlation, λ, the degree to which this trait evolution deviates from Brownian motion ^38^, was evaluated by fitting PGLS models with different values of λ to find that which maximized the log-likelihood of the best-fitted model. The level of statistical significance was set at alpha = 0.05.

### Phylogenetic analysis

This was performed by PGLS on the 203 species’ specific geographical location, habitat, ΔT and number of individuals measured. As fishes’ physiology is dependent on the environmental thermal conditions, hemisphere was incorporated into the model because of the significant differences in thermal variability between the two hemispheres ^6^, with the north having higher thermal variation than the south ^27^. Due to the effects of local thermal variation on fish thermal physiology, we included an interaction term between latitudinal groups (tropical versus temperate) and the ΔT. We also conducted general linear model (GLM) analysis to exclude the effect of phylogeny on the outcome of the observed variation in log_10_ S.D.CT_max_, testing the individual effects of our variables in the model (suppl. Table 3).

## Authors contributions

J.J.H. Nati, M.B.S. Svendsen, S. Marras, S.S. Killen, J.F. Steffensen, D.J. McKenzie, P. Domenici designed the study. J.J.H. Nati and S.S. Killen performed the statistical analyses. J.J.H. Nati wrote the manuscript. J.J.H. Nati, M.B.S. Svendsen, S. Marras, S.S. Killen, J.F. Steffensen, D.J. McKenzie, P. Domenici revised the manuscript.

