## Supplementary material for "Intraspecific variation in thermal tolerance differs between tropical and temperate fishes"

**Supplementary Methods and Results**

1) **Model selection approach:** to test differences in intraspecific variation in $CT_{max}$ between two relevant latitudinal groups, habitat types, hemisphere effect and methodology effects.

PGLS models from the most complex to the simple one:

Model 1 : $\log_{10}$ S.D. $CT_{max}$ ~ $\alpha_0 + \alpha_1$ Latitudinal position + $\alpha_2$ Habitat + $\alpha_3$delta Temperature+ $\alpha_4$ Hemisphere + $\alpha_5$ Number of individuals used + $\alpha_6$ Latitudinal position x delta Temperature + $\varepsilon$

Model 2 : $\log_{10}$ S.D. $CT_{max}$ ~ $\alpha_0 + \alpha_1$ Latitudinal position + $\alpha_2$ Habitat + $\alpha_3$delta Temperature + $\alpha_4$ Hemisphere + $\alpha_5$ Number of individuals used + $\varepsilon$

Model 3 : $\log_{10}$ S.D. $CT_{max}$ ~ $\alpha_0 + \alpha_1$ Latitudinal position + $\alpha_2$ Habitat + $\alpha_3$delta Temperature + $\alpha_4$ Hemisphere + $\varepsilon$

Model 4 : $\log_{10}$ S.D. $CT_{max}$ ~ $\alpha_0 + \alpha_1$ Latitudinal position + $\alpha_2$ Habitat + $\alpha_3$delta Temperature + $\varepsilon$

Model 5 : $\log_{10}$ S.D. $CT_{max}$ ~ $\alpha_0 + \alpha_1$ Latitudinal position + $\alpha_2$ Habitat + $\varepsilon$

Model 6 : $\log_{10}$ S.D. $CT_{max}$ ~ $\alpha_0 + \alpha_1$ Latitudinal position + $\varepsilon$

Model 7 : $\log_{10}$ S.D. $CT_{max}$ ~ $\alpha_0 + \varepsilon$

Supplementary Table 1: PGLS model selection approach by AIC function on 203 fish species. Selected model is model 1 due to lowest AIC value.

| N species | models | df | AIC |
|---|---|---|---|
| 203 | Model 1 | 7 | 268.5 |
| | Model 2 | 6 | 277.3 |
| | Model 3 | 5 | 279.8 |
| | Model 4 | 4 | 313 |
| | Model 5 | 3 | 327.7 |
| | Model 6 | 2 | 330.2 |
| | Model 7 | 1 | 344.5 |

### 2) Phylogenetic informed analysis on intraspecific variation of $CT_{max}$ in 203 species

Model 1 : $\log_{10}$ S.D. $CT_{max}$ ~ $\alpha_0$ + $\alpha_1$ Latitudinal position + $\alpha_2$ Habitat + $\alpha_3$delta Temperature + $\alpha_4$ Hemisphere + $\alpha_5$ Number of individuals used + $\alpha_6$ Latitudinal position x delta Temperature + ε

Supplementary Table 2: PGLS model summary, $F_{6,195} = 4.397$, $\lambda = 0.553$, $R^2 = 11.92$, $p < 0.001$ on 203 fish species.

| N species | coefficients | estimates | s.e. | t values | p values |
|---|---|---|---|---|---|
| 203 | Intercept | -0.393 | 0.184 | -2.137 | 0.034 |
| | Tropical species | -0.51 | 0.248 | -2.054 | 0.041 |
| | Marine species | -0.115 | 0.068 | -1.683 | 0.094 |
| | Delta Temperature | 0.014 | 0.007 | 1.972 | 0.05 |
| | Southern Hemisphere | 0.151 | 0.065 | 2.318 | 0.022 |
| | Individuals | 0.002 | 0.002 | 0.846 | 0.399 |
| | Tropical sp*delta Temperature | 0.038 | 0.019 | 1.95 | 0.053 |

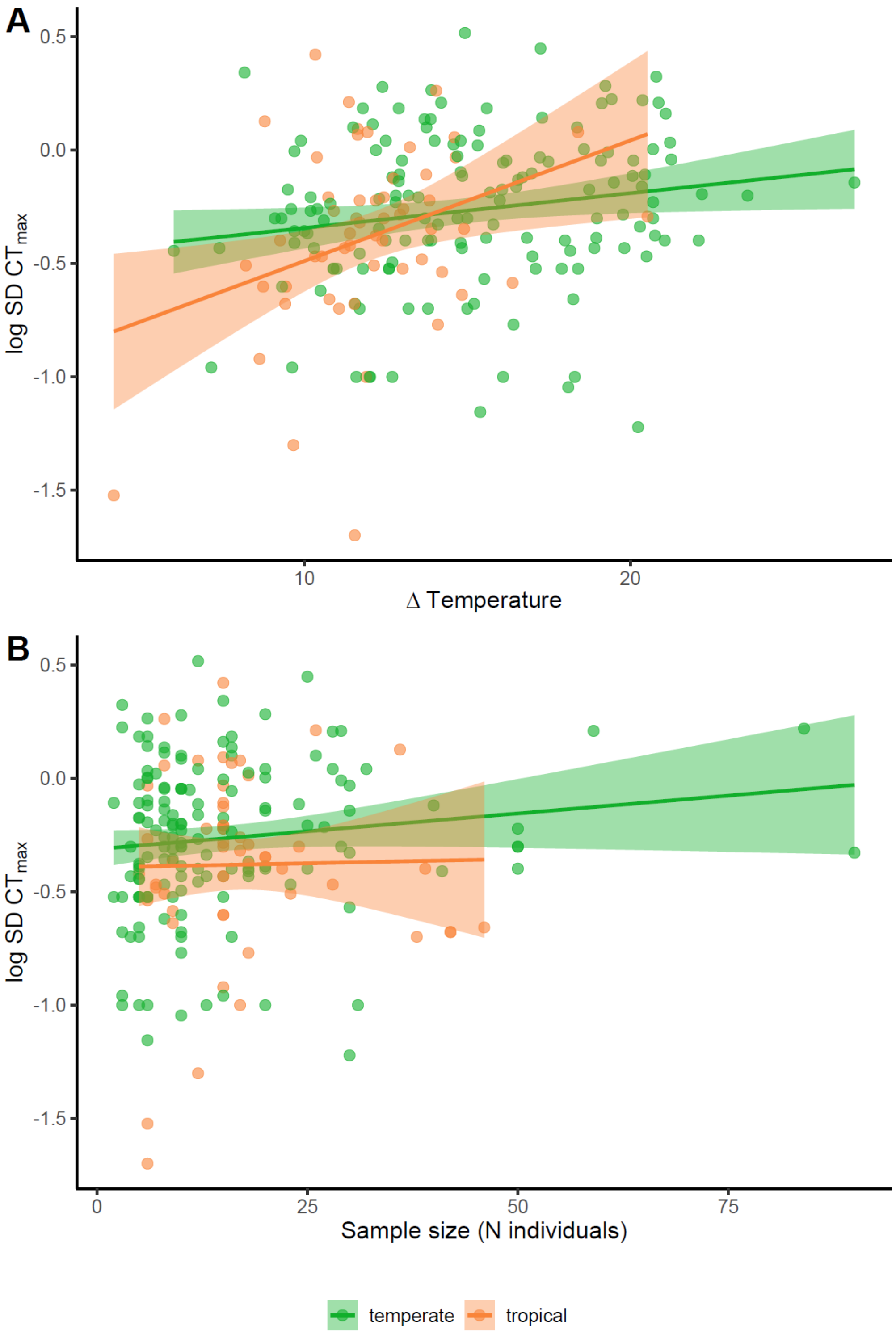


**Figure S1| The effect of (A) delta temperature and (B) the number of individuals used in each study on $log_{10}$ transformed standard deviation of $CT_{max}$ ($log_{10}$ S.D. $CT_{max}$) divided by two latitudinal groups (tropical and temperate species).** The shaded areas in the regression lines correspond to 95% of confidence interval.

### 3) GLM analysis on intraspecific variation $CT_{max}$ in 203 species

Supplementary Table 3: GLM model summary on $log_{10}$ S.D.$CT_{max}$, $F_{6,196} = 4.24$ , $R^2 = 11.49$, $p < 0.001$ in 203 fish species.

| N species | coefficients | estimates | s.e. | t values | p values |
|---|---|---|---|---|---|
| 203 | Intercept | -0.483 | 0.115 | -4.210 | < 0.001 |
| | Tropical species | -0.529 | 0.26 | -2.038 | 0.043 |
| | Marine species | -0.095 | 0.053 | -1.790 | 0.075 |
| | Delta Temperature | 0.013 | 0.007 | 1.751 | 0.082 |
| | Southern Hemisphere | 0.123 | 0.067 | 1.842 | 0.067 |
| | Individuals | 0.002 | 0.002 | 1.283 | 0.201 |
| | Tropical sp*Delta Temperature | 0.037 | 0.02 | 1.818 | 0.071 |

### 4) Supplementary Figures

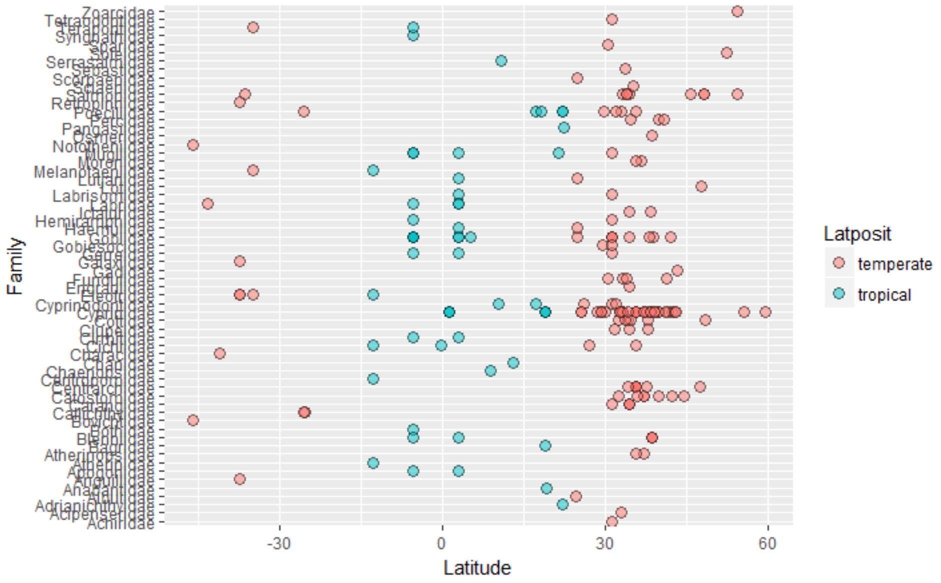


**Figure S2| Representation of family across latitude divided in two latitudinal groups.** red dot temperate species, blue dots in tropical species.

### 5) Phylogenetic tree

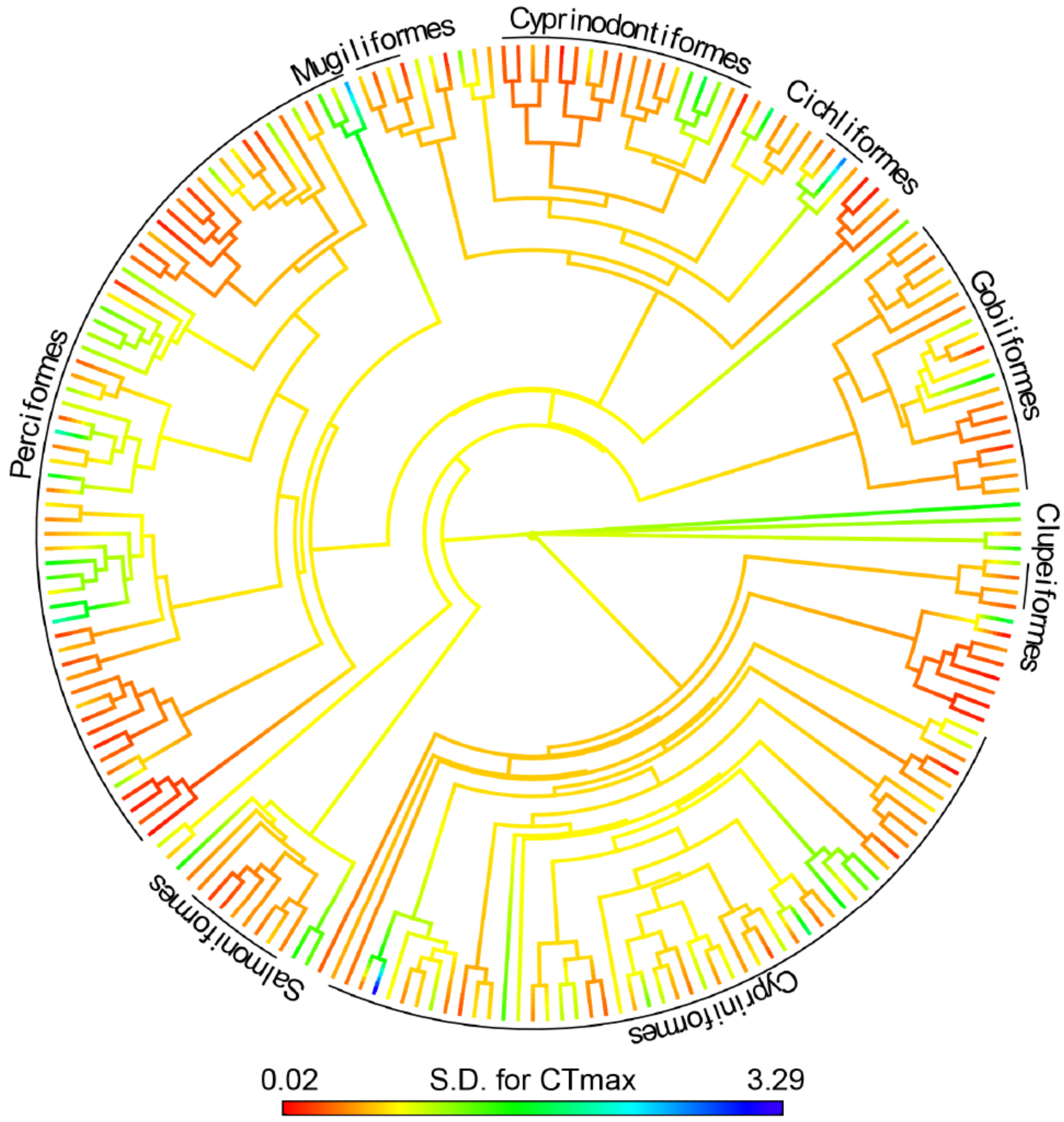


**Figure S3| Phylogenetic tree of 203 species and their families, organised according to their intraspecific variation in upper thermal tolerance, estimated as the standard deviation of their $CT_{max}$ (S.D. for $CT_{max}$).**
